# Co-occurrence and cooperation between comammox and anammox bacteria in a full-scale attached growth municipal wastewater treatment process

**DOI:** 10.1101/2022.12.05.519185

**Authors:** Katherine Vilardi, Irmarie Cotto, Megan Bachmann, Mike Parsons, Stephanie Klaus, Christopher Wilson, Charles Bott, Kelsey Pieper, Ameet Pinto

## Abstract

Cooperation between comammox and anammox bacteria for nitrogen removal has been recently reported in laboratory-scale systems including synthetic community construct; however, there are no reports of full-scale municipal wastewater treatment systems with such cooperation. Here, we report intrinsic and extant kinetics as well as genome-resolved community characterization of a full-scale integrated fixed film activated sludge (IFAS) system where comammox and anammox bacteria co-occur and appear to drive nitrogen loss. Intrinsic batch kinetic assays indicated that majority of the aerobic ammonia oxidation was driven by comammox bacteria (1.75 ± 0.08 mg-N/g TS-h) in the attached growth phase with minimal contribution by ammonia oxidizing bacteria. Interestingly, a portion of total inorganic nitrogen (∼8%) was consistently lost during these aerobic assays. Aerobic nitrite oxidation assays eliminated the possibility of denitrification as a cause of nitrogen loss, while anaerobic ammonia oxidation assays resulted in rates consistent with anammox stoichiometry. Full-scale experiments at different dissolved oxygen (DO = 2-6 mg/L) set points indicated persistent nitrogen loss that was partly sensitive to DO concentrations. Genome-resolved metagenomics confirmed high abundance (relative abundance 6.53 ± 0.34%) of two *Brocadia-*like anammox populations while comammox bacteria within the *Ca*. Nitrospira nitrosa cluster were lower in abundance (0.37% ± 0.03%) and *Nitrosomonas*-like ammonia oxidizers even lower (0.12% ± 0.02%). Collectively, our study reports for the first time the co-occurrence and co-operation of comammox and anammox bacteria in a full-scale municipal wastewater treatment system.

**Synopsis:** Comammox and anammox cooperation resulted in dissolved oxygen concentration dependent nitrogen loss in municipal wastewater treatment system.

## INTRODUCTION

Despite their ubiquitous detection in engineered and natural ecosystems^1–7^, the role of comammox bacteria in full-scale nitrogen removal remains to be established. Our previous work demonstrated that comammox bacteria are most prevalent in nitrogen removal systems treating wastewater with an attached growth phase or long solids retention times, and they often co-occur with strict AOB and *Nitrospira*-NOB^8^. Further, a large portion of comammox bacteria detected in wastewater systems, including those in our past studies^8,9^, belong to clade A1 comammox bacteria and are affiliated with *Ca*. Nitrospira nitrosa-like populations^10–12^. However, kinetic parameters for the majority of comammox bacteria are undetermined; only one isolated species (*Ca*. Nitrospira inopinata)^13^ and one enrichment (*Ca*. Nitrospira krefti)^14^ have demonstrated a high affinity for ammonia. Assessment of ammonia oxidation activity in wastewater treatment systems with coexisting strict AOB and comammox bacteria has been done using metatranscriptomics which suggested comammox bacteria were active and potentially metabolically flexible^15^. However, quantifying nitrification rates of comammox bacteria in wastewater treatment systems would help better define their roles in nitrogen removal from wastewater and their ecological niche relative to other nitrifying bacteria.

Recent literature has demonstrated the potential for comammox bacteria to cooperate with anammox bacteria for efficient nitrogen conversion to dinitrogen gas in laboratory-scale systems^16– 19^. This cooperation between comammox and anammox bacteria could take on different modalities. For instance, comammox bacteria could provide nitrite to anammox bacteria through partial nitrification of ammonia^16,17^ or comammox bacteria could perform complete nitrification to nitrate which is then converted to nitrite by denitrifying bacteria for use by anammox bacteria^18^. Both modalities involving comammox-anammox bacterial co-operation could be potentially more beneficial as compared to traditional strategies involving ammonia oxidizing bacteria (AOB) as this would minimize the potential for biotic nitrous oxide (N_2_O) production^20^. Comammox bacteria have a lower affinity for ammonia compared to strict AOB^13,14^ which could be important in ammonia-limited environments. Further, suppression of nitrite oxidizers is necessary to ensure anammox bacteria do not washout of the system^6^. Thus, comammox bacteria could limit nitrite availability to strict NOB under aerobic conditions if they perform complete nitrification to nitrate. Studies have demonstrated comammox bacteria associated partial nitrification-anammox achieved 70% nitrogen removal under low DO conditions with suspended biomass^17^, while systems with a biofilm phase demonstrated nitrogen removal under both high^16^ and low^18^ DO conditions. Further, Cui et al (2022) reported the enrichment of *Ca*. Nitrospira nitrosa-like comammox bacteria in a predominantly anammox system when operated under microaerobic conditions. Though some studies suggest comammox bacteria prefer oxygen limited conditions^6,18^, our previous survey of comammox bacteria in different wastewater treatment systems did not find any association between the prevalence/abundance of comammox bacteria and DO concentrations^8^. To date, all four studies reporting comammox-anammox co-occurrence and cooperation for nitrogen removal are laboratory-scale systems. There are currently no reports of comammox-anammox cooperation in mainstream full-scale nitrogen removal wastewater systems.

In this study, we report the co-occurrence and cooperation of comammox and anammox bacteria for nitrogen removal in a full-scale integrated fixed film activated sludge (IFAS) system. Our previous studies at the Hampton Roads Sanitation District (HRSD) James River Treatment Plant (JRTP) in Virginia, USA found high abundance of comammox bacteria in the attached growth phase with their concentration routinely exceeding those of canonical AOB^8,9^. Follow-up experiments to determine the nitrification kinetics of comammox bacteria indicated the potential for the presence of anammox bacteria including nitrogen loss consistent with anammox stoichiometry. Thus, we systematically characterized the intrinsic and extant (i.e., *in situ*) kinetics of aerobic and anaerobic ammonia removal and the microbial community using genome-resolved metagenomics to identify the nitrifying populations responsible for nitrogen removal at full-scale.

## MATERIALS AND METHODS

### Treatment plant description

A full-scale 20 million gallons per day (MGD) integrated fixed film activated sludge (IFAS) system for treating municipal wastewater (average COD = 400-500 mg/L, average ammonia = 20-40 mg/L) was monitored in December 2021. The sampled IFAS system was one of nine parallel treatment trains that operate in an A2O configuration, including R1 (anaerobic), R2-R3 (anoxic), R4 (aerobic), and R5 (deaeration) (Figure 1A). The total volumes of the anaerobic, anoxic, and aerobic zones per train are 0.06, 0.13, and 0.26 MG, respectively. The aerobic zone contains plastic carrier media (AnoxKaldnes K3, specific surface area 500 m^2^/m^3^) with attached biomass growth (Figure 1) at a percent carrier fill of 45%. The plastic carrier media are kept suspended by coarse bubble aeration. Two internal mixed liquor recycle (IMLR) pumps transfer nitrate to the anoxic zones, one with suction from one side of the upstream portion of R4 and one from the downstream end of R4.

**Figure 1:**
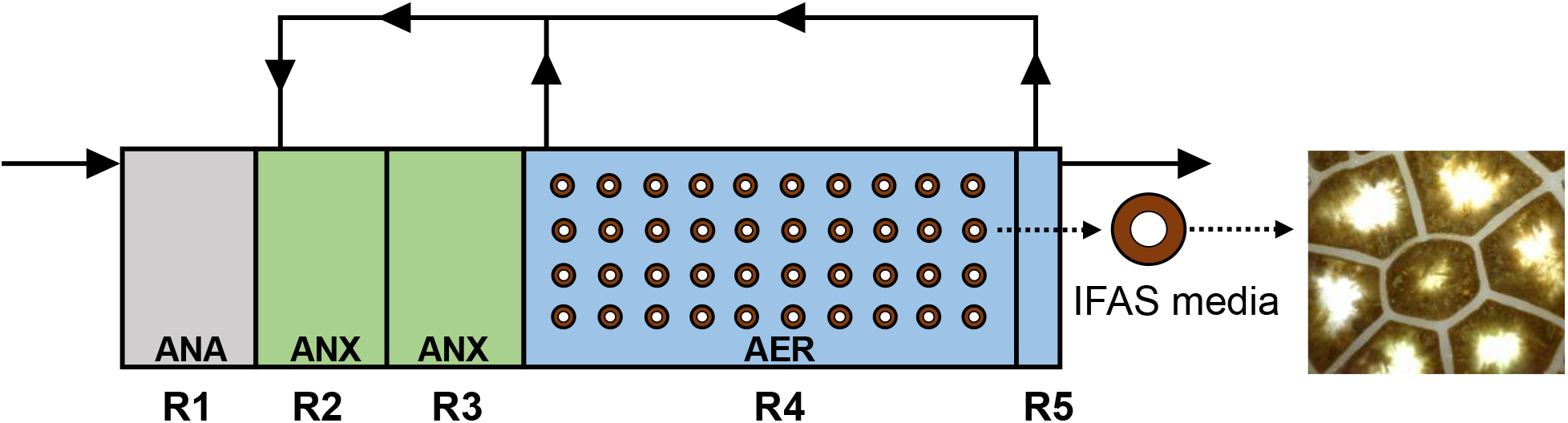
Schematic of full-scale IFAS system and image of IFAS biofilm taken with Dino-Lite Digital using the Dino-Lite 2.0 Software microscope. ANA = Anaerobic, ANX = Anoxic, AER = Aerobic.

### Intrinsic kinetics assays for evaluating aerobic ammonia and nitrite oxidation, and anaerobic ammonium oxidation rates

Media and suspended solids were collected from the aerobic IFAS zone for intrinsic kinetics assays (Section SI-1). All assays for the attached phase were carried out using 40 pieces of media in 900 mL of secondary clarifier effluent. Assays for the suspended phase used 900 mL of mixed liquor. Mixing was maintained in beakers for both phases with a magnetic st1irbar. Aerobic ammonia and nitrite oxidation batch assays were performed with a 28 mg-N/L initial concentration of NH_4_^+^-N (by spiking ammonium chloride stock solution) and NO_2_^-^-N (by spiking in sodium nitrite stock solution), respectively, while tests for anaerobic ammonium oxidation were performed without aeration and 28 mg-N/L spikes of both NH_4_^+^-N and NO_2_^-^-N. Differential inhibition batch assays were conducted with 28 mg NH_4_^+^-N/L and spikes of either 4 μM 1-octyne to inhibit AOB^21,22^ or 100 μM allylthiourea (ATU)^23,24^ to inhibit all aerobic ammonia oxidation. Inhibitors were added to beakers 30 minutes before the nitrogen spikes. Dissolved oxygen (DO), pH, and temperature were measured before the nitrogen spike and at the end of each assay using the Thermo Scientific Orion Star A329 Portable pH/ISE/Conductivity/RDO/DO meter (Cat. No. STARA3290).

Aqueous samples were collected and filtered through a 0.45-μm syringe filter (Sartorius, Cat. No. 14-555-278) immediately after spiking in nitrogen and every 15 minutes for the subsequent hour. The total inorganic nitrogen was calculated by adding the measured concentrations of ammonia, nitrite, and nitrate as nitrogen at each timepoint. The change in ammonia, nitrite, nitrate, and total inorganic nitrogen concentrations over time were then used to obtain corresponding rates (mg-N/L-hr). These rates were converted to specific rates (mg-N/g TS-h) by dividing with the average concentration of total attached biomass or total suspended solids (Section SI-2). To compare comammox and anammox ammonia oxidation rates with those reported in literature, abundance adjusted rates (μmol N/mg protein-h) were calculated by dividing the average ammonia consumption rate (mg-N/g TS-h) obtained from aerobic or anaerobic ammonia oxidation batch assays by the portion of total metagenomic reads mapping to comammox or anammox bacteria metagenome assembled genomes (see below) as their approximate contribution to total solids measured and then using the conversion factor 1.9 mg dry weight/mg protein^25^.

### Full-scale experiments with variable dissolved oxygen (DO) setpoints

Full-scale experiments were conducted by varying the DO concentration of the aerobic IFAS zone to four setpoints: 2, 3, 4, and 6 mg/L on four consecutive days. Reactor monitoring and sample characterization details are provided in supplementary material (Section SI-2). Duplicate samples were collected across the full-scale A2O system for each setpoint approximately six hours after adjustment to the new DO setpoint (Section SI-3). *In situ* rates of ammonia oxidation, nitrate production, and loss of total inorganic nitrogen in the aerobic zone were calculated using measurements obtained from samples collected at the end of the anoxic zone (influent to aerobic zone) and at the end of the aerobic zone) (Section SI-4).

### Metagenomic sequencing and data processing

Biomass attached to six pieces of media collected from the aeration tank were scrapped using a sterile scalpel and homogenized using a sterile loop. 250 mg of biomass from each sample was then used for DNA extraction using Qiagen’s DNeasy Powersoil Pro Kit (Cat. No. 47016) on the Qiacube (Cat. No. 9002160). DNA concentrations were measured using Invitrogen Qubit dsDNA Broad Range Kit (Cat. No. Q32850). DNA extracts were subject to library preparation using NEBNext Ultra II FS DNA Library Prep Kit followed by sequencing on the Illumina NovaSeq 6000 platform in 2×250 bp mode on a single SP Flowcell by the Molecular Evolution Core at the Parker H. Petit Institute for Bioengineering and Bioscience at Georgia Institute of Technology.

Raw short reads were trimmed to remove low quality bases/reads using fastp v0.22.0^26^, and the Univec database was used to remove contamination from the filtered reads (Table SI-2). Clean reads from Sample2 were assembled into contigs using metaSpades v3.15.5^27^ with kmer sizes of 21, 33, 55, and 77. The resulting fasta files were indexed with bwa index v0.7.17^28^, and the paired end reads were mapped to the assembly using bwa mem v0.7.17. The resulting sam files were converted to bam files using ‘samtools view -F 4 -bhS’ using SAMtools v1.15.1^29^ to retain only mapped reads. We did not perform a co-assembly with reads from the six IFAS pieces to avoid increasing the complexity of the sample. In general, wastewater samples exhibit high diversity and, even at high coverage, co-assemblies can be extremely challenging. Thus, while pooling samples reads would increase the genome coverage, increasing the sample complexity may result in fewer assembled genomes.

Binning was performed using MetaBAT2 v2.15^30^, CONCOCT v1.1.0^31^, and MaxBin2 v2.2.7^32^ with contigs greater than 2000 bp with DAStool v1.1.4^33^ used to combine and curate the refined bins to generate a non-redundant set of bins. The quality and taxonomy of the resulting bins were determined with CheckM v1.2.1^34^ and the Genome Taxonomy Database Toolkit (GTDB-Tk 2.1.1, database release r207 v2)^35,36^, respectively. The assembly and bins were subject to gene calling using Prodigal v2.6.3^37^ and gene annotation against the KEGG database^38^ using kofamscan v1.3.0^39^ to explore the genes associated with aerobic and anaerobic ammonia oxidation and nitrite oxidation (i.e., *amoA* [KO number K10944], *amoB* [K10945], *amoC* [K10946], *hao* [K10535], *nxrA* [K00370], *nxrB* [K00371], *hzs* [K20932], *hdh* [K20935]).

*Brocadia* (n=2) and *Nitrospira* (n=3) MAGs recovered from this study (Table SI-3) were phylogenetically placed in context of 90 and 85 previously publicly available *Brocadia* and *Nitrospira* genomes, respectively, using Anvi’o v7.1^40^. The *Nitrospira* references included nine previously assembled *Nitrospira* MAGs (i.e., 7 NOB and 2 comammox MAGs) from samples taken in 2017-2018 from the same system. All other reference genomes were obtained from NCBI (Table SI-4). ORFs were predicted using Prodigal v2.6.3 and then searched against a collection of HMM models (Bacteria_71) including 38 ribosomal proteins, summarized by Lee (2019)^41^ using hmmscan v3.2.^42^. Multiple pairwise alignments for each gene were performed using MUSCLE v3.8.1551^43^. Each phylogenomic tree was constructed using ITOL v2.1.7^44^. No comammox bacteria or AOB (e.g., *Nitrosomonas*) MAGs were assembled from this sample. Therefore, the *amoA* gene sequences found in the metagenomic assembly were aligned using BLAST with the comammox (n=2) and *Nitrosomonas* (n=9) MAGs previously obtained from the same IFAS system (Cotto 2022) to verify whether these populations were still present. Maximum likelihood phylogenetic tree of *Nitrospira*-comammox and *Nitrosomonas*, based on the *amoA* gene, were performed aligning them with MUSCLE v3.8.1551 followed by construction of the tree using IQ-TREE v2.0.3^45^.

In order to calculate the relative abundances of the nitrifying bacteria, MAGs generated from this and the past study^9^ were grouped and dereplicated using drep v2.5.4^46^ at 95% ANI with completeness and contamination thresholds set to 50% and 10%, respectively. Only MAGs with genome coverage (proportion of the genome covered by at least one read) higher than 50% in each sample were used to calculate relative abundances. The genome coverage and relative abundance of each MAG in reads per kilobase million (RPKM) per sample were calculated with coverM v0.6.1 (https://github.com/wwood/CoverM). The percent relative abundance of each MAG was calculated by mapping all sample reads to each genome and dividing the resulting mapped reads by the total reads in that sample. All sequencing data along with MAGs are deposited in NCBI under BioProject number PRJNA908221.

## RESULTS AND DISCUSSION

### Comammox bacteria are the principal active aerobic ammonia oxidizers in the attached growth phase

Aerobic intrinsic kinetics assays indicated that the specific ammonia oxidation rate (sAOR) was 2.5 times lower for the suspended solids (0.694 mg-N/g TS-h) (Figure SI-1) compared to the attached phase (1.75 ± 0.08 mg-N/g TS-h) (Figure 2A). This indicates that approximately 71% of the ammonia oxidation capacity was in the biofilm which is consistent with prior work at JRTP^47^ and Broomfield Wastewater Treatment^48^. However, the intrinsic kinetic rates measured in this study were significantly lower than previously reported from the same system^47^. Specifically, the average specific NOx production rate (sNPR) measured in this study were 1.22 ± 0.09 mg-N/g

**Figure 2:**
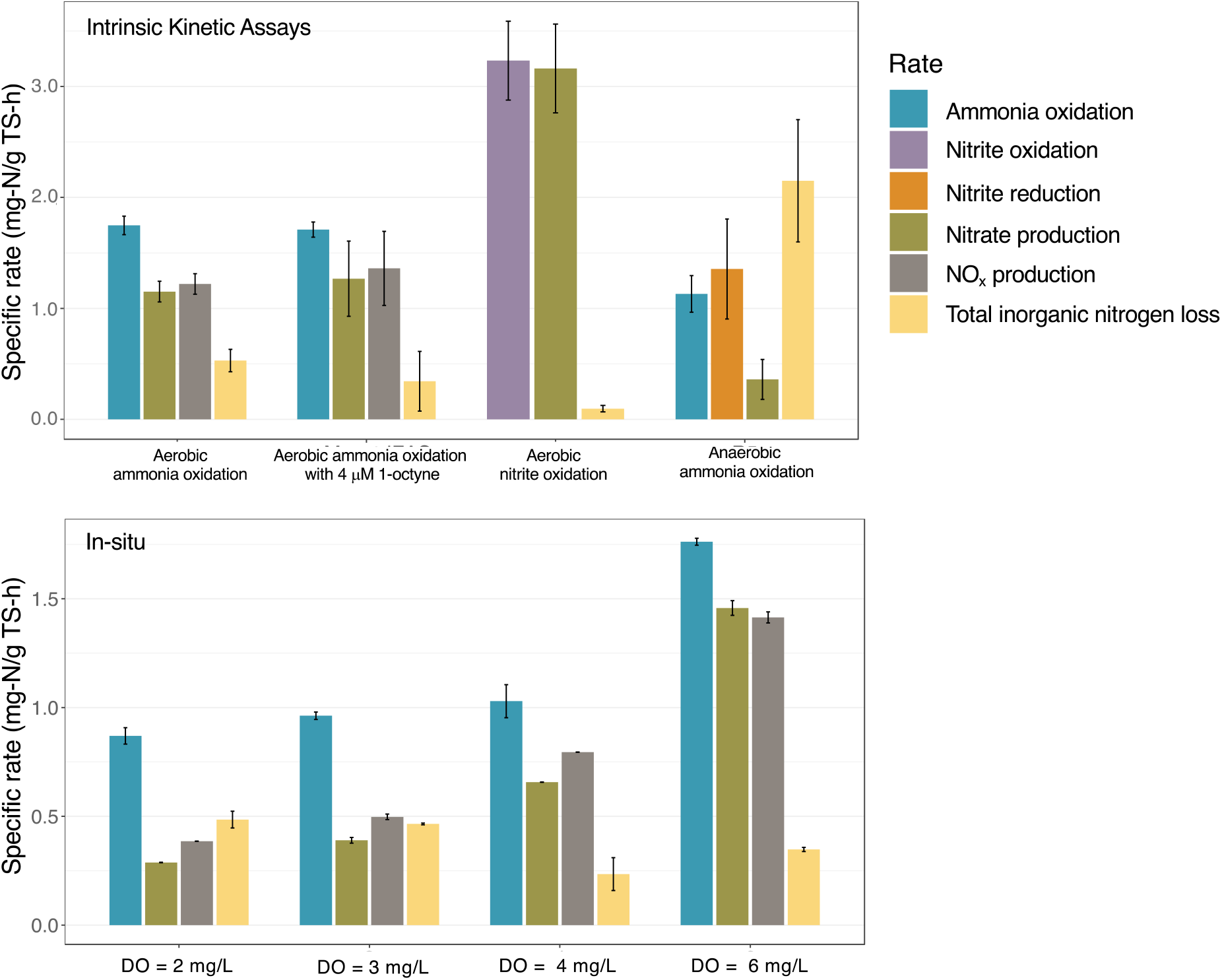
(A) Intrinsic rates of ammonia oxidation, nitrite plus nitrate production (NO_X_), nitrate production, and total inorganic nitrogen loss for aerobic (uninhibited and inhibited with 4 μM 1-octyne.) and anaerobic ammonia oxidation and aerobic nitrite oxidation assays. (B) Rates of ammonia oxidation, nitrate production, and total inorganic nitrogen loss in the aerobic zone at DO set points estimated for full-scale experiment. Error bars denote variation across replicate batch assays (Figure 2A) and replicate measurements in full-scale system (Figure 3B).

TS-h (Figure 2A, Table SI-1) as compared to 2.39-5.87 mg-N/g TS-h in previous work using IFAS media^47,48^. The differing rates between systems could be a result of differences in nitrifier community composition, as well as methods used to determine total solids in the attached phase.

Addition of ATU in the kinetic assay with suspended solids resulted in complete cessation of ammonia oxidation whereas ammonia oxidation and corresponding NOx production occurred in the assay spiked with 1-octyne comparable to uninhibited assay. Addition of ATU also resulted in the complete cessation of aerobic ammonia oxidation with no nitrite or nitrate accumulation. However, with 4 μM 1-octyne, the average sAOR was 1.69 ± 0.06 mg-N/g TS-h, indicating aerobic ammonia oxidation was not substantially inhibited (Figure 2A) (p>0.05, unpaired t-test). This occurred despite irreversible inhibition of strict AOB reported at 1-octyne concentrations as low as 1 μM^21^ with no inhibition of either ammonia oxidizing archaea (AOA) or comammox bacteria^22^. Our prior study suggested comammox bacteria were the dominant aerobic ammonia oxidizer in the attached phase, while strict AOB were comparatively lower in abundance^8^. Thus, taken together, these results suggest that comammox bacteria were likely the principal aerobic ammonia oxidizer in the attached phase. Considering this, we focused our remaining work on the attached phase microbial community.

### Loss of total inorganic nitrogen occurs in both aerobic and anaerobic ammonia oxidation conditions

Interestingly, we observed substantial total inorganic nitrogen loss (∼8%) during both, the uninhibited and 1-octyne spiked ammonia oxidation assays (TIN loss rate: 0.57 ± 0.09 mg-N/g TS-h (Figure 2A) for attached phase biomass assays. This loss was likely not due to denitrification since the DO concentrations in the aerobic batch assays were maintained at 6 mg/L. To confirm this, we performed aerobic nitrite oxidation assays under identical conditions as aerobic ammonia oxidation assays which revealed a nearly closed nitrogen balance (0.6 - 1.2% gap in nitrogen balance) (Figure 2A). This prompted us to investigate anaerobic ammonia oxidation as a possibility mode of nitrogen loss in attached phase assays. Anaerobic ammonium oxidation assays revealed a total inorganic nitrogen loss rate greater than nitrate produced (Figure 2A) (TIN loss rate: 2.15 ± 0.55 g-N/g TS-h). Further, the proportion of the ammonia to nitrite consumption rate (1:1.20), ammonium consumption to nitrate production rate (1:0.32), and rate of nitrogen loss (1.84) were indicative of anammox bacterial activity^49^.

The capacity for both aerobic and anaerobic ammonia oxidation has been observed in low DO (∼0.5 mg/L) bench-scale demonstrations established from wastewater^17,18^. However, we observed a loss of total inorganic nitrogen under both aerobic (6 mg/L) and anaerobic ammonia oxidization conditions, suggesting that anammox activity may not be completely inhibited by higher DO conditions. This could potentially be due to anammox bacteria existing in oxygen-limited parts of the IFAS biofilm and nitrite made available by aerobic ammonia oxidation used by anammox bacteria to drive a loss of nitrogen^16^. Though nitric oxide (NO) and nitrous oxide (N_2_O) were not measured in batch assays as possible forms of nitrogen loss, stoichiometric evidence strongly supports loss of total inorganic nitrogen was due to anammox bacteria in the attached phase. While strict AOB can produce N_2_O via NO through nitrifier denitrification, differential inhibition assays indicated that comammox bacteria were the primary aerobic ammonia oxidizers in the IFAS media. It has been demonstrated that at least one comammox bacteria species (i.e., *Ca*. Nitrospira inopinata) cannot denitrify to N_2_O and produces N_2_O comparable to AOA which is substantially lower than that of AOB^20^. Therefore, it is unlikely that nitrogen loss was due to N_2_O production.

### Dissolved oxygen dependent nitrification and nitrogen loss occur in the full-scale IFAS system

Since anammox activity was observed in batch assays, experiments were performed to quantify this activity *in situ* in the full-scale system by modifying the DO concentration of the aerobic zone to set points ranging from 2-6 mg/L and monitoring process parameters (Figure 2C, Section SI-3, Figure SI-2 and 3). Interestingly, total inorganic nitrogen loss (∼10%) was still observed despite the high DO set point (Figure 2C, Figure SI-4). However, the two lowest set points, 2 and 3 mg/L had higher rates of total inorganic nitrogen loss than nitrate production (16% nitrogen loss) (Figure SI-4). Thus, in these instances, anaerobic activity mediated by anammox bacteria was enhanced compared to higher DO settings (Figure SI-5). It is important to note that some of the nitrogen loss and its estimation could be attributable to assimilation and/or ammonification of organic nitrogen.

The highest ammonia oxidation rate (1.76 mg-N/g TS-h) was obtained at a DO setpoint of 6 mg/L; this rate was similar to what was observed in the batch assays. Ammonia oxidation rates at DO set points 2, 3, and 4 mg/L were similar to each other (0.870-1.03 mg/g TS-h) and were 42-51% lower compared to the rate observed at 6 mg/L. Comparatively, the percent ammonia removed was 31, 35, 41 and 63% at DO set points 2, 3, 4 and 6 mg/L, respectively (Figure SI-4). The sharp decrease in ammonia oxidation rates with lowering of DO setpoints could be due to oxygen limitation within the biofilm^47,50^. However, Zhao et al (2022)^51^ recently demonstrated a similar dramatic decrease in ammonia oxidation rates in a comammox enrichment moving bed biofilm reactor dominated by two *Ca*. Nitrospira nitrosa-like populations. Specifically, they report a 50% decrease in ammonia oxidation rate with decrease in DO concentrations from 6 to 2 mg/L and attribute this to low apparent oxygen affinity of *Ca*. Nitrospira nitrosa-like bacteria (K_o_=2.8 mg O_2_/L). This would appear consistent with our observations in the full-scale system, further suggesting that comammox bacteria were the primary drivers of aerobic ammonia oxidation.

In this study, the portion of *in situ* ammonia oxidized aerobically by comammox bacteria increased with DO while the portion oxidized by anammox bacteria was higher at lower DO conditions (Figure 3) indicating that lower DO concentrations reduce the aerobic ammonia oxidation rate of comammox bacteria while simultaneously favoring conditions for anaerobic ammonia oxidation. Further, this demonstrates comammox and anammox bacteria can cooperate at a low enough DO such that comammox bacteria can still make nitrite available for anammox bacteria who in turn can drive a loss of total inorganic nitrogen. The implications of operating at much lower DO, such as those suggested in other studies^6,18^ (less than 1 mg/L), may limit comammox bacterial ammonia oxidation such that they are unable to produce nitrite for anammox bacteria.

**Figure 3:**
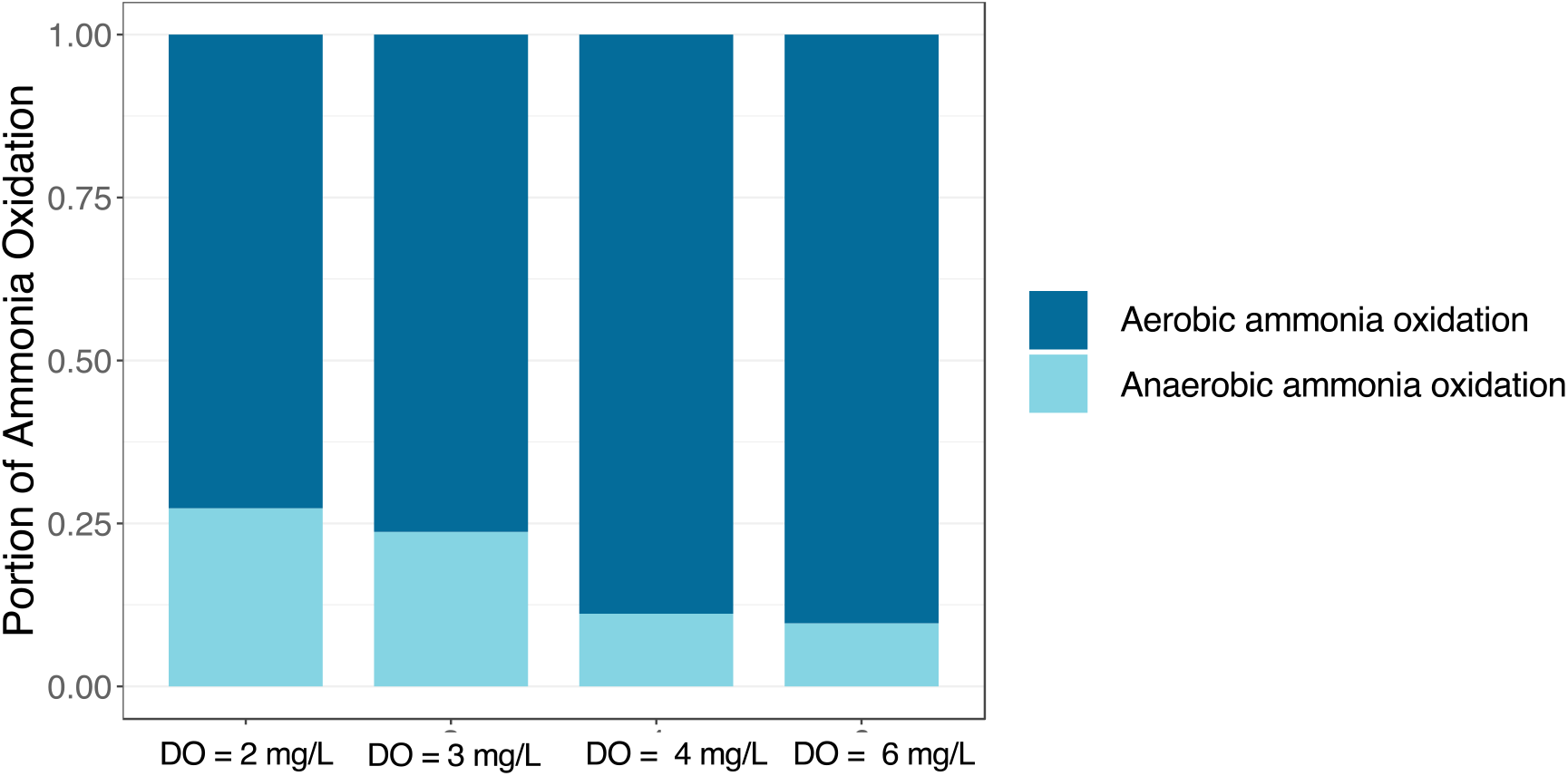
Portion of the total ammonia oxidation rate attributed to aerobic and anaerobic ammonia oxidation.

### Low abundance comammox bacteria co-occur with highly abundant anammox bacteria in IFAS media

No comammox or strict AOB MAGs were assembled from samples collected during this study which is in contrast to our previous assembly of two comammox and nine *Nitrosomonas* MAGs from this IFAS system^9^. Two *amoA* gene sequences in the metagenomic assembly were aligned using BLAST with previously assembled MAGs obtained from the same IFAS system. These *amoA* sequences showed a greater than 99 and 97% sequence identity, respectively with the *amoA* genes present in one comammox and one *Nitrosomonas* MAGs previously assembled^9^ (Figure 4A). Further, contigs obtained from the metagenomic assembly in this study were aligned with BLAST against previously assembled nitrifier MAGs associated with comammox bacteria, *Nitrospira*-NOB, and *Nitrosomonas*. This revealed that several contigs in this study were fully aligned (zero mismatches, 100% ID) to these previously nitrifier MAGs suggesting that the comammox bacteria and AOB were present in the samples, but at very low abundances and thus their genomes were not successfully reconstructed.

**Figure 4:**
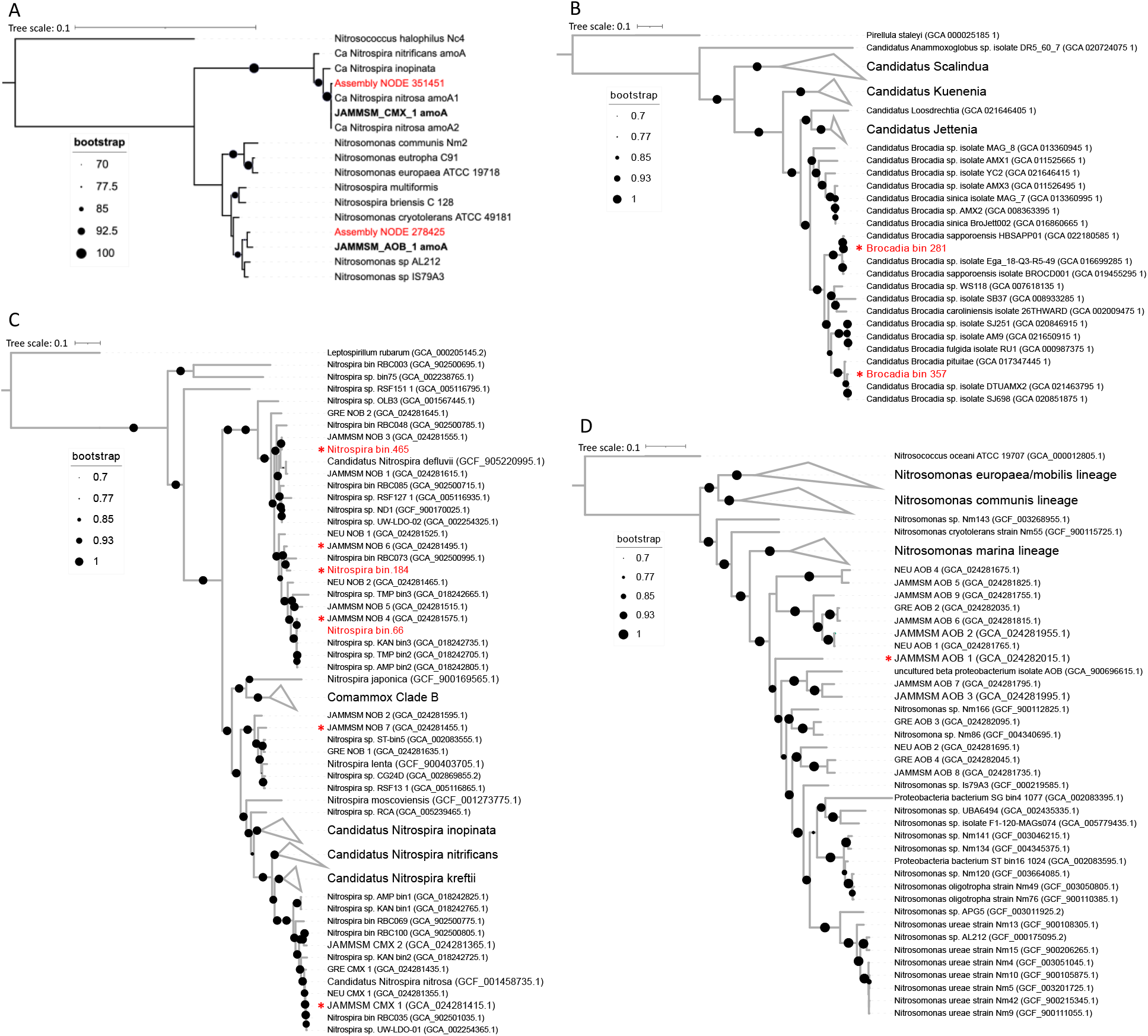
(A) Maximum likelihood phylogenetic tree of the *amoA* genes found in the assembly (red) along with *Nitrospira*-comammox and *Nitrosomonas amoA* genes references (black). Black references in bold are *amoA* sequences in MAGs recovered from our previous study (2017-2018). Phylogenetic placement of (B) *Brocadia*, (C) *Nitrospira* and (D) *Nitrosomonas* MAGs with 90, 85 and 65 reference genomes, respectively. Branches that are not related to any relevant MAG are collapsed. The complete list of the reference genomes used in the analysis is in Table SI-4. Red labels are MAGs recovered from this study, black labels are genome references downloaded from NCBI and black bold labels are MAGs from samples taken from 2017 to 2018 in the same IFAS system. MAGs selected after dereplication and used to calculate the relative abundances of nitrifying bacteria are marked with a red asterisk.

MAGs associated with *Brocadia* (n=2) and *Nitrospira* (n=3) were obtained from biomass attached to IFAS media even though anammox bacteria were not found in our past study^8,9^. *Nitrospira* and *Brocadia* MAGs represented 6.53 ± 0.34 % and 6.25 ± 1.33% of total reads in the sample. Phylogenomic analysis associated *Brocadia*-like MAGs with *Ca*. Brocadia sapporoensis and *Ca*.

Brocadia pituitae (Figure 4B) while *Nitrospira*-like MAGs were all placed in lineage I associated with *Nitrospira defluvii* (Figura 4C). Two *Nitrospira* MAGs were very similar to three *Nitrospira*-lineage I MAGs assembled from samples taken between 2017 and 2018 in the same IFAS system (Cotto 2022) (ANI = 99.95% for Nitrospira_bin.66 and JAMMSM_NOB_4, 99.29% for Nitrospira_bin.465 and JAMMSM_NOB_3 and, 96.40% for Nitrospira_bin.465 and JAMMSM_NOB_1). However, these *Nitrospira* MAGs were at much lower abundances in this study (8.35 ± 1.91 RPKM) compared with the previous study (55.56 ± 14.71 RPKM)^9^.

The decrease in the *Nitrospira* abundance could be the reason why several of the previously assembled MAGs could not be assembled in the current study despite the fact that 5 out of 7 of the previously assembled *Nitrospira* MAGs had 90% of their genomes covered using reads from this study (Figure 5A). Therefore, the relative abundance of all nitrifying groups was calculated from a set of dereplicated MAGs recovered from both studies (Table SI-3). However, only MAGs with genome coverage (i.e., percent of the genome covered by reads from this study) higher than 50% (Table SI-5) were selected for relative abundance calculations (Table SI-6). The results confirm the presence of most previously assembled *Nitrospira* MAGs (including one *Nitrospira* lineage II MAG) in the system but at much lower abundances (17.26 ± 3.61 RPKKM) compared with the samples from 2017-2018 (122.90 ± 30.69 RPKM) (Figure 5B, Table SI-6). Further, the genome coverage of previously assembled comammox (JAMMSM_CMX_1) and *Nitrosomonas* (JAMMSM_AOB_1) MAGs were 80.6 ± 9.8 and 72.3 ± 1.0%, respectively (Table SI-5). In conjunction with the *amoA* (Figure 4A) and contig level analysis, this confirms the presence of previously detected comammox and *Nitrosomonas* genomes in the system. Thus, relative abundance estimates of comammox bacteria and *Nitrosomonas* were calculated by mapping the reads from the six IFAS samples to these previously assembled MAGs (Figure 4C and D).

**Figure 5:**
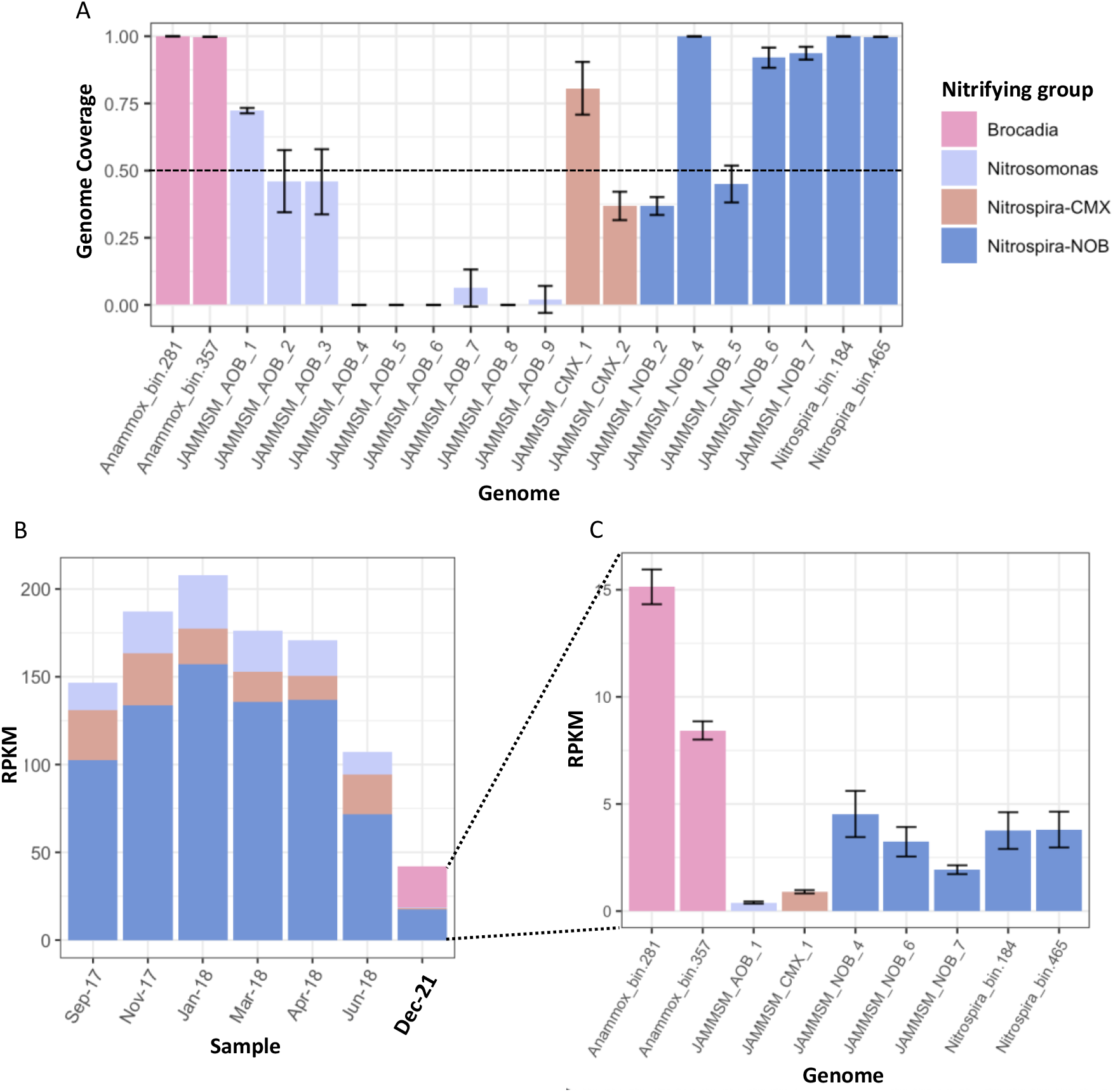
(A) Average genome coverage of dereplicated MAGs in six samples (IFAS media pieces) taken from the aeration tank. Error bars represent the standard deviation across the six samples. MAGs with genome coverage higher than 50% were considered present in the system and used to calculate the relative abundances of the nitrifying groups (i.e., *Brocadia, Nitrosomonas, Nitrospira*-comammox, and *Nitrospira*-NOB). (B) Cumulative relative abundances in reads per kilobase million (RPKM) of *Brocadia, Nitrosomonas, Nitrospira*-comammox (CMX) and *Nitrospira*-NOB obtained from the current study samples (December-2021) and samples taken between September 2017 and June 2018. (C) Average relative abundance of each genome in the IFAS pieces taken on December-2021. Error bars represent the standard deviation across the six samples.

Comammox and *Nitrosomonas* relative abundances were about 0.90 ± 0.8 RPKM and 0.40 ± 0.05 RPKM, respectively (Figure 5C). This differs from our prior work, where comammox and *Nitrosomonas* relative abundances were 22 ± 6.26 and 21.04 ± 6.17 RPKM, respectively (Figure 5B). Thus, it is very likely that the low abundance of comammox bacteria and *Nitrosomonas* affected the assembly and binning process, which did not allow for the reconstruction of these genomes even though they are still present in the system. Despite the decrease in both comammox and *Nitrosomonas* relative abundance in the system, the comammox:*Nitrosomonas* proportion is higher in this study relative to our previous work in the same system^8,9^. These results, coupled with the inhibition kinetic assays with 1-octyne and drop in in situ ammonia oxidation rate with decrease in DO suggests that comammox bacteria are the principal aerobic ammonia oxidizers in this system. The abundance adjusted ammonia consumption rate for comammox bacteria was 64.19 μmol-N/mg protein-h which is within the range reported for isolated *Ca. Nitrospira inopinata* (14 μmol-N/mg protein-h)^13^ and enriched *Ca. Nitrospira kreftii* (83 μmol-N/mg protein-h)^14^. Additionally, the adjusted rate for anammox bacteria was 2.37 μmol-N/mg protein-h which is similar to the reported rate for other anammox bacteria (3.27 μmol-N/mg protein-h)^52^. Anammox bacteria outnumbered comammox bacteria and strict AOB despite high bulk DO of the IFAS system favoring aerobic ammonia oxidizers. While recent studies have suggested that anammox bacteria are most likely oxygen tolerant rather than strictly anaerobic^53,54^, the comparatively high abundance of anammox in the attached phase could also be due to anaerobic zones deeper in the biofilm. Further, transcriptional activity of anammox genes associated with *Brocadia* were found in aquifers with anoxic-to-oxic conditions suggesting anammox bacteria are able to contribute to nitrogen loss in a diverse range of oxygen environments^55^.

### Co-operative nitrogen removal by comammox and anammox bacteria

Comammox-anammox co-occurrence has been previously demonstrated in synthetic community constructs and/or lab-scale reactors using attached growth phases^16,18,19^. Spatial organization as a contributor to comammox-anammox cooperation was highlighted by Gotshall 2020 where comammox bacteria form a protective outer layer where oxygen was most available while anammox bacteria occupy inner biofilm layers. Cooperation could also be aided by their differing affinities for nitrite since comammox bacteria have a lower affinity for nitrite than anammox bacteria^13,52^. At JRTP, nitrite made available from ammonia oxidation by comammox bacteria was used by anammox bacteria along with residual ammonia to generate a loss of total inorganic nitrogen. In this IFAS system, influent to the aerobic zone contained limited nitrite (Figure SI-3). Therefore, comammox-driven ammonia oxidization was likely the primary source of nitrite production in the aerobic zone, which occurs predominantly in the attached phase and not in the suspended phase. One potential reason for a comparably lower sAOR/sNPR could be explained by the low abundance and slower nitrification rates of comammox bacteria. For example, Onnis-Hayden 2007 estimated the relative abundance of their nitrifying community was about 10% and 15-20% *Nitrosomonas*-like ammonia oxidizers and *Nitrospira*-like bacteria, respectively, with sNPR rates approximately three times higher than the rates observed in this study. Our full-scale results show loss of total inorganic nitrogen at various DO concentrations, suggesting anammox bacteria were shielded from complete DO inhibition in aerobic environments. However, aerobic nitrification was still the dominant process at each tested DO set point (SI Fig-5). While nitrate accumulation under full-scale conditions demonstrates that strict NOB or comammox bacteria used majority of the produced nitrite, the estimated rates suggest that anammox bacteria used a portion of it to drive a loss of total inorganic nitrogen at each tested DO concentration. Nitrite affinities (K_s_) for *Nitrospira*-NOB and anammox bacteria associated with MAGs in this study are similar (*Nitrospira defluvii* (9 μM)^56^, and *Ca*. Brocadia sapporoensis (5 μM)^57^) while the reported value for the one isolated comammox bacteria *Nitrospira* inopinata (449.2 μM)^13^ is much lower. Thus, *Nitrospira*-NOB and anammox bacteria may outcompete comammox bacteria for nitrite, and the decrease in nitrogen loss with increase in ammonia oxidation and nitrate production rates at higher DO concentrations suggests the competition was oxygen dependent.

To our knowledge, this is the first report of a full-scale main-stream system with cooccurring comammox-anammox populations. Here, we show the potential for cooperation between comammox and anammox bacteria for mainstream systems across a range of DO concentration. Our results suggests that DO-dependent reduction in the ammonia oxidation rate of comammox bacteria maximizes nitrogen loss via anammox activity, while higher DO concentrations result in nitrate accumulation not only due to lower anammox rates but due to higher ammonia oxidation rates of comammox bacteria.

## Supporting information

- Methodological details, additional figures, and tables are provided in Supplemental Materials.

## Acknowledgement

This research was supported by NSF CBET 1703089 and NSF CBET 1923124.

