## Supplementary Material for "Co-occurrence and cooperation between comammox and anammox bacteria in a full-scale attached growth municipal wastewater treatment process"

### DESCRIPTION OF SUPPORTING INFORMATION

Section S1. Batch assay set-up and experiments

Section S2. Solids Measurements

Section S3. Sample collection and processing for full-scale experiments

Section S4. In situ rate calculations

Table S1: Rates obtained from triplicate aerobic ammonia and nitrite oxidation, and anaerobic ammonia oxidation batch assays

Table S2: Total number of raw reads and reads remaining in sample fastq files after quality filtering with fastp

Figure S1: Specific rates for ammonia oxidation, nitrate production, and loss of total inorganic nitrogen for the suspended solids.

Figure S2: Chemical and physical process parameters measured along the transect of the IFAS system.

Figure S3: Nitrogen species as a percent of influent total inorganic nitrogen along the transect of the IFAS system.

Figure S4: Percent of ammonia and total inorganic nitrogen removal in the aerobic zone at each DO set point.

Figure S5: Specific rates for aerobic and anaerobic nitrogen biotransformation in the aerobic zone of the full-scale IFAS system during DO set point experiments

**Section S1. Preparation of Batch Assay Set-up and Experiments**

Stock solutions were prepared as 10,000 mg $NH_4$-N/L (using ammonium chloride) and 10,000 mg $NO_2$-N/L (using sodium nitrite). Prior to conducting experiments, the IFAS media was collected from the aerobic zone and washed overnight in secondary clarifier effluent to remove loosely associated biomass. 1 L glass beakers were set up on stir plates with a magnetic bar placed inside along with an aeration stone connected to an aquarium pump to provide aeration for aerobic assays. After washing with secondary effluent, 40 pieces of IFAS media were added to the beakers with 900 mL of secondary clarifier effluent for the attached phase assays while 900 mL mixed liquor were added to beakers for suspended phase assays. The aquarium pump was turned on to provide aeration for aerobic experiments with pump settings modified based on DO measurements to achieve a DO concentration between 6-8 mg/L. Alkalinity and pH in all beakers were measured before the experiments and adjusted using the sodium bicarbonate stock solution to ensure adequate alkalinity for nitrification. Blank controls were established for each condition using 900 mL of filtered water.

**Section S2. Sample characterization and total solids measurements**

DO concentrations and temperature were measured directly at each sampling location in the full-scale IFAS system using a probe (IPM insiteIG with PortaCaddie) submerged in the wastewater. pH was measured using the HACH Company Pocket Pro+ pH meter (Cat. No. 9532000) immediately after sample collection. Duplicate samples of mixed liquor were filtered at all locations in the field through 0.45 μM syringe filters for chemical analysis. Hach Company TNT Vials were used to determine concentrations of ammonia (TNT832, TNT831), nitrite (TNT839, TNT840), nitrate (TNT835), chemical oxygen demand (TNT821, TNT820), and total alkalinity (TNT870) of the filtered aqueous samples from both full-scale and intrinsic kinetic experiments. All samples were analyzed on a HACH DR3900 photospectrometer (Cat. No. LPV440.99.00002). IFAS media and suspended solids from both ends of the aerobic zone were collected on the sampling days to measure total solids (TS) and total suspended solids (TSS). Suspended solids and IFAS media were also collected from both ends of the aerobic zone for DNA extraction for microbial characterization.

Three IFAS pieces and suspended solids samples were taken from two aeration zones (beginning and end) on three different sampling dates. The suspended solids concentration (g/L) was measured using Standard Method 2540D. All IFAS pieces (n=18) were baked overnight at 105 °C and then weighed before scrubbing off all biomass with a 2 N $H_2SO_4$ solution and small bristle brush. The clean IFAS pieces were dried overnight at 105 °C and then weighed. The mass of total solids (TS) on individual pieces were obtained by subtracting the mass of the cleaned IFAS piece from the mass recorded for dried attached biomass. The average mass of TS per media piece for each sampling date was then multiplied by 40 (number of IFAS pieces used in each assay) to estimate the total mass of attached biomass in batch assays for the attached phase. The average concentration of total attached biomass in grams per liter was estimated by dividing the average total mass of attached biomass (g) by 0.9 L (volume of secondary clarifier effluent used in batch assays). Total attached biomass in the aerobic zone of the treatment train was estimated by multiplying the average mass of attached biomass obtained from all 18 pieces by the number of IFAS pieces in the aerobic zone (~114155600). The average concentration of total attached biomass (g/L) was estimated by dividing the aforementioned average (g) by the tank volume, 984207 L.

**Section S3. Sample collection and processing for full-scale experiments**

The six sampling locations selected included the anoxic zone (n=3, one per stage for R1, R2, and R3), aerobic zone (n=2, one in the beginning (R4_Z1) and one at the end (R4_Z4)), and deaeration (n=1, R5) (Figure 1 main manuscript). Samples were collected using a long handle scooper approximately six hours after the DO concentration was modified in the aerobic zone, and only one DO setting was applied per day. In the field, IFAS media was stored by cutting pieces with a sterile razor blade and placed two per tube into 15 mL falcon tubes for DNA extraction. Both IFAS media and suspended solids samples for DNA extraction were placed in a -20 °C freezer at the treatment plant until they were shipped on dry ice to Northeastern University, where they remained in a -80 °C freezer until extraction. Samples were taken at five designated sampling locations along the transect on days when the DO was set to 2, 4, and 6 mg/L for measuring volatile fatty acids (VFAs). Seven VFAs, including acetic, butyric, caproic, isobutyric, isovaleric, propionic and valeric acid, were surveyed using standard method SM5560D, but only acetic acid was detected above the limit of quantification (LOQ = 5 mg/L) in any of the wastewater samples.

**Section S4. In situ rate calculations**

In situ rates of ammonia oxidation, nitrite oxidation, nitrate production, and loss of total inorganic nitrogen in the aerobic zone were calculated using the following equations:

Ammonia oxidation rate ($rNH_3$) = (($NH_{3,in}$ - $NH_{3,eff}$)/Aerobic HRT)/TS (1)

Nitrate production rate ($rNO_3$) = (($NO_{3,eff}$ - $NO_{3,in}$)/Aerobic HRT)/TS (2)

Total inorganic nitrogen removal rate (rTIN) = ((($NH_{3,in}$ + $NO_{2,in}$ + $NO_{3,in}$) - ($NH_{3,eff}$ + $NO_{2,eff}$ + $NO_{3,eff}$))/ Aerobic HRT)/TS (3)

Anaerobic ammonia oxidation rate ($rNH_{3,ANA}$) = rTIN/2.04 (4)

Anaerobic nitrite reduction rate ($rNO_{2,ANA}$) = 1.32(rTIN/2.04) (5)

Anaerobic nitrate production rate ($rNO_{3,ANA}$ ) = 0.26(rTIN/2.04) (6)

Aerobic ammonia oxidation ($rNH_{3,AER}$) = $rNH_3$ - $rNH_{3,ANA}$ (7)

Aerobic nitrate production rate ($rNO_{3,AER}$) = $rNO_3$ - $rNO_{3,ANA}$ (8)

Average concentrations (mg-N/L) for ammonia, nitrite, and nitrate were obtained from duplicates samples collected from the influent to the aerobic zone (subscript “in”) at the final anoxic zone at R3 and effluent from the aerobic zone (subscript “eff”) at R4_Z4. Total inorganic nitrogen (mg/L) was determined by the summation of ammonia, nitrite, and nitrate concentrations measured at R3 and R4_Z4. The hydraulic retention time (HRT) of the aerobic zone expressed in hours was calculated by dividing the volume of the aerobic tank by the flowrate across the aerobic zone on day of sampling. Estimates for anaerobic ammonia oxidation ($rNH_{3,ANA}$), nitrite reduction ($rNO_{2,ANA}$) and nitrate production ($rNO_{3,ANA}$) rates were calculated using the constants derived from anammox reaction stoichiometry corresponding to 2.04 (1 mol $1NH_4^+$ consumed produces 1.02 mol $N_2$: $1NH4^+ + 1.32NO_2 \rightarrow 1.02N_2 + 0.26NO_3^-$) of nitrogen (as N), 0.26 mol of nitrate (as N) and 1.32 mol of nitrite (as N) per mole of ammonium consumed. All rates are expressed as specific rates (mg-N/g TS-hr) calculated by dividing the *in situ* rates by the estimated concentration of total attached biomass in the aerobic zone.

**Table S1**: Rates obtained from triplicate aerobic ammonia and nitrite oxidation, and anaerobic ammonia oxidation batch assays for the attached growth phase.

| **Assay** | **Rate type** | **Average rate (mg-N/g TS-h)** | **Standard deviation** |
|---|---|---|---|
| Aerobic ammonia oxidation | Ammonia removal | 1.75 | 0.08 |
| | Nitrate production | 1.15 | 0.09 |
| | NOx production | 1.22 | 0.09 |
| | TIN loss | 0.53 | 0.10 |
| Aerobic nitrite oxidation | Nitrite oxidation | 3.23 | 0.36 |
| | Nitrate production | 3.16 | 0.40 |
| | TIN loss | 0.10 | 0.03 |
| Anaerobic ammonia oxidation | Ammonia removal | 1.13 | 0.17 |
| | Nitrite reduction | 1.36 | 0.45 |
| | Nitrate production | 0.36 | 0.18 |
| | TIN loss | 2.15 | 0.55 |

TIN: total inorganic nitrogen

**Table S2**: Total number of raw reads and reads remaining in sample fastq files after quality filtering with fastp

| Sample | Raw | Filtered |
|---|---|---|
| 1 | 251564616 | 243864532 |
| 2 | 209058090 | 202372028 |
| 3 | 208199202 | 201819776 |
| 4 | 220372610 | 212874004 |
| 5 | 192953414 | 186805330 |
| 6 | 198420028 | 191168754 |

**Table S3**: Set of nitrifier MAGs recovered from samples taken from June 2017 to June 2018 and one sample (Sample2) from December 2021 after dereplication.

| **Genome** | **Completeness** | **Redundancy** | **Length** | **Contigs** |
|---|---|---|---|---|
| Brocadia _bin.281* | 92.00 | 0.00 | 2868031 | 125.00 |
| Brocadia_bin.357* | 90.00 | 0.00 | 2601165 | 241.00 |
| JAMMSM_AOB_1 | 94.12 | 9.80 | 2926721 | 166.00 |
| JAMMSM_AOB_2 | 100.00 | 2.63 | 2859940 | 22.00 |
| JAMMSM_AOB_3 | 100.00 | 0.00 | 2882584 | 21.00 |
| JAMMSM_AOB_4 | 96.08 | 0.00 | 1957621 | 82.00 |
| JAMMSM_AOB_5 | 98.04 | 5.26 | 2568442 | 19.00 |
| JAMMSM_AOB_6 | 92.16 | 1.96 | 2031342 | 65.00 |
| JAMMSM_AOB_7 | 94.12 | 2.63 | 3066737 | 73.00 |
| JAMMSM_AOB_8 | 98.04 | 2.63 | 2970458 | 91.00 |
| JAMMSM_AOB_9 | 78.43 | 1.96 | 2167635 | 21.00 |
| JAMMSM_CMX_1 | 94.12 | 1.96 | 4065392 | 23.00 |
| JAMMSM_CMX_2 | 86.27 | 1.96 | 2629561 | 220.00 |
| JAMMSM_NOB_2 | 90.20 | 0.00 | 2828782 | 50.00 |
| JAMMSM_NOB_4 | 100.00 | 0.00 | 3829071 | 5.00 |
| JAMMSM_NOB_5 | 70.59 | 2.63 | 2923168 | 59.00 |
| JAMMSM_NOB_6 | 88.24 | 7.89 | 2856077 | 119.00 |
| JAMMSM_NOB_7 | 84.31 | 0.00 | 2879078 | 61.00 |
| Nitrospira_bin.184* | 94.00 | 6.00 | 4812459 | 231.00 |
| Nitrospira_bin.465* | 90.00 | 2.00 | 3222716 | 235.00 |

*MAGs recovered from this study

**Table S4**: Publicly available *Nitrospira*, *Brocadia* and *Nitrosomonas* genomes downloaded from NCBI. Genomes previously assembled from JRTP by Cotto et al 2022 are shown in bold.

| **Reference genomes** | **Accession number** |
|---|---|
| Leptospirillum rubarum | GCA_000205145.2 |
| Candidatus Nitrospira defluvii | GCA_000196815.1 |
| Candidatus Nitrospira inopinata | GCF_001458695.1 |
| Candidatus Nitrospira kreftii isolate | GCA_014058405.1 |
| Candidatus Nitrospira nitrificans | GCF_001458775.1 |
| Candidatus Nitrospira nitrosa | GCF_001458735.1 |
| Nitrospira bin RBC001 | GCA_902500755.1 |
| Nitrospira bin RBC003 | GCA_902500695.1 |
| Nitrospira bin RBC035 | GCA_902501035.1 |
| Nitrospira bin RBC042 | GCA_902500795.1 |
| Nitrospira bin RBC044 | GCA_902500705.1 |
| Nitrospira bin RBC048 | GCA_902500785.1 |
| Nitrospira bin RBC069 | GCA_902500775.1 |
| Nitrospira bin RBC073 | GCA_902500995.1 |
| Nitrospira bin RBC083 | GCA_902500745.1 |
| Nitrospira bin RBC085 | GCA_902500715.1 |
| Nitrospira bin RBC093 | GCA_902500825.1 |
| Nitrospira bin RBC100 | GCA_902500805.1 |
| Nitrospira japonica | GCF_900169565.1 |
| Nitrospira lenta | GCF_900403705.1 |
| Nitrospira moscoviensis | GCF_001273775.1 |
| Nitrospira sp. AMP bin1 | GCA_018242825.1 |
| Nitrospira sp. AMP bin2 | GCA_018242805.1 |
| Nitrospira sp. bin75 | GCA_002238765.1 |
| Nitrospira sp. CG24A | GCA_002869925.2 |
| Nitrospira sp. CG24B | GCA_002869845.2 |
| Nitrospira sp. CG24C | GCA_002869885.2 |
| Nitrospira sp. CG24D | GCA_002869855.2 |
| Nitrospira sp. CG24E | GCA_002869895.2 |
| Nitrospira sp. Ga0074138 | GCA_001464735.1 |
| Nitrospira sp. KAN bin1 | GCA_018242765.1 |
| Nitrospira sp. KAN bin2 | GCA_018242725.1 |
| Nitrospira sp. KAN bin3 | GCA_018242735.1 |
| Nitrospira sp. LK265 | GCA_011090395.1 |
| Nitrospira sp. LK70 | GCA_011090425.1 |
| Nitrospira sp. MAG 137 | GCA_009594955.1 |

| | |
|---|---|
| Nitrospira sp. MAG 149 | GCA_009594935.1 |
| Nitrospira sp. MAG 237 | GCA_009595005.1 |
| Nitrospira sp. MAG 242 | GCA_009594945.1 |
| Nitrospira sp. MAG 245 | GCA_009594825.1 |
| Nitrospira sp. MAG 246 | GCA_009594995.1 |
| Nitrospira sp. MAG 248 | GCA_009594835.1 |
| Nitrospira sp. MAG 266 | GCA_009594865.1 |
| Nitrospira sp. MAG 275 | GCA_009594855.1 |
| Nitrospira sp. MAG 79 | GCA_902500715.1 |
| Nitrospira sp. ND1 | GCF_900170025.1 |
| Nitrospira sp. OLB3 | GCA_001567445.1 |
| Nitrospira sp. RCA | GCA_005239465.1 |
| Nitrospira sp. RCB | GCA_005239475.1 |
| Nitrospira sp. RSF1 1 | GCA_005116965.1 |
| Nitrospira sp. RSF11 1 | GCA_005116945.1 |
| Nitrospira sp. RSF12 1 | GCA_005116955.1 |
| Nitrospira sp. RSF127 1 | GCA_005116935.1 |
| Nitrospira sp. RSF13 1 | GCA_005116865.1 |
| Nitrospira sp. RSF151 1 | GCA_005116795.1 |
| Nitrospira sp. RSF3 1 | GCA_005116835.1 |
| Nitrospira sp. RSF48 1 | GCA_005116775.1 |
| Nitrospira sp. RSF5 1 | GCA_005116895.1 |
| Nitrospira sp. RSF6 1 | GCA_005116885.1 |
| Nitrospira sp. RSF7 1 | GCA_005116825.1 |
| Nitrospira sp. RSF8 1 | GCA_005116815.1 |
| Nitrospira sp. RSF9 1 | GCA_005116745.1 |
| Nitrospira sp. SCGC | GCA_001644405.1 |
| Nitrospira sp. SCN 59-13 | GCA_001724505.1 |
| Nitrospira sp. SG-bin1 | GCA_002083365.1 |
| Nitrospira sp. SG-bin2 | GCA_002083405.1 |
| Nitrospira sp. ST-bin4 | GCA_002083565.1 |
| Nitrospira sp. ST-bin5 | GCA_002083555.1 |
| Nitrospira sp. TMP bin1 | GCA_018242685.1 |
| Nitrospira sp. TMP bin2 | GCA_018242705.1 |
| Nitrospira sp. TMP bin3 | GCA_018242665.1 |
| Nitrospira sp. UW-LDO-01 | GCA_002254365.1 |
| Nitrospira sp. UW-LDO-02 | GCA_002254325.1 |
| Nitrospira sp. WS110 | GCA_011090365.1 |
| GRE_NOB_1 | GCA_024281635.1 |

| | |
|---|---|
| GRE_NOB_2 | GCA_024281645.1 |
| NEU_NOB_1 | GCA_024281525.1 |
| NEU_NOB_2 | GCA_024281465.1 |
| **JAMMSM_NOB_1** | **GCA_024281615.1** |
| **JAMMSM_NOB_2** | **GCA_024281595.1** |
| **JAMMSM_NOB_3** | **GCA_024281555.1** |
| **JAMMSM_NOB_4** | **GCA_024281575.1** |
| **JAMMSM_NOB_5** | **GCA_024281515.1** |
| **JAMMSM_NOB_6** | **GCA_024281495.1** |
| **JAMMSM_NOB_7** | **GCA_024281455.1** |
| Nitrosococcus oceani ATCC 19707 | GCA_000012805.1 |
| Nitrosomonas aestuarii strain Nm36 Ga0181080 101 | GCF_003046585.1 |
| Nitrosomonas aestuarii strain Nm69 | GCF_900114305.1 |
| Nitrosomonas communis strain Nm110 | GCF_900106545.1 |
| Nitrosomonas communis strain Nm2 chromosome | GCF_001007935.1 |
| Nitrosomonas communis strain Nm44 | GCF_900114745.1 |
| Nitrosomonas cryotolerans strain Nm55 | GCF_900115725.1 |
| Nitrosomonas europaea isolate OLB2 | GCA_001567435.1 |
| Nitrosomonas europaea strain ATCC 25978 | GCF_900167395.1 |
| Nitrosomonas eutropha C91 | GCF_000014765.1 |
| Nitrosomonas eutropha strain Nm19 Ga0181055 101 | GCF_003046355.1 |
| Nitrosomonas eutropha strain Nm24 | GCF_900116685.1 |
| Nitrosomonas eutropha strain Nm38 | GCF_900106875.1 |
| Nitrosomonas halophila strain Nm1 | GCF_900107165.1 |
| Nitrosomonas marina strain Nm22 | GCF_900110145.1 |
| Nitrosomonas marina strain Nm71 | GCF_900111605.1 |
| Nitrosomonas mobilis isolate 1 | GCF_900103035.1 |
| Nitrosomonas nitrosa strain Nm90 Ga0181054 101 | GCF_003051105.1 |
| Nitrosomonas oligotropha strain Nm49 Ga0181079 101 | GCF_003050805.1 |
| Nitrosomonas oligotropha strain Nm76 | GCF_900110385.1 |
| Nitrosomonas sp. AL212 | GCF_000175095.2 |
| Nitrosomonas sp. APG5 | GCF_003011925.2 |
| Nitrosomonas sp. Is79A3 | GCF_000219585.1 |
| Nitrosomonas sp. isolate F1-120-MAGs074 | GCA_005779435.1 |
| Nitrosomonas sp. isolate UBA8640 | GCA_003514765.1 |
| Nitrosomonas sp. Nm120 Ga0180021 11 | GCF_003664085.1 |
| Nitrosomonas sp. Nm132 | GCF_900100485.1 |

| | |
|---|---|
| Nitrosomonas sp. Nm134 Ga0181067 101 | GCF_004345375.1 |
| Nitrosomonas sp. Nm141 Ga0181066 101 | GCF_003046215.1 |
| Nitrosomonas sp. Nm143 Ga0207393 101 | GCF_003268955.1 |
| Nitrosomonas sp. Nm166 | GCF_900112825.1 |
| Nitrosomonas sp. Nm33 | GCF_900107265.1 |
| Nitrosomonas sp. Nm34 | GCF_900113925.1 |
| Nitrosomonas sp. Nm51 | GCF_900111165.1 |
| Nitrosomonas sp. Nm58 | GCF_900107335.1 |
| Nitrosomonas sp. Nm86 Ga0181061 101 | GCF_004340695.1 |
| Nitrosomonas sp. UBA6494 | GCA_002435335.1 |
| Nitrosomonas stercoris KYUHI-S | GCA_006742785.1 |
| Nitrosomonas ureae strain Nm10 | GCF_900105875.1 |
| Nitrosomonas ureae strain Nm13 | GCF_900108305.1 |
| Nitrosomonas ureae strain Nm15 | GCF_900206265.1 |
| Nitrosomonas ureae strain Nm42 | GCF_900215345.1 |
| Nitrosomonas ureae strain Nm4 Ga0181063 1001 | GCF_003051045.1 |
| Nitrosomonas ureae strain Nm5 Ga0181075 101 | GCF_003201725.1 |
| Nitrosomonas ureae strain Nm9 | GCF_900111055.1 |
| Proteobacteria bacterium SG bin4 1077 | GCA_002083395.1 |
| Proteobacteria bacterium ST bin16 1024 | GCA_002083595.1 |
| uncultured beta proteobacterium isolate AOB | GCA_900696615.1 |
| GRE_AOB_1 | GCA_024282195.1 |
| GRE_AOB_2 | GCA_024282035.1 |
| GRE_AOB_3 | GCA_024282095.1 |
| GRE_AOB_4 | GCA_024282045.1 |
| NEU_AOB_1 | GCA_024281765.1 |
| NEU_AOB_2 | GCA_024281695.1 |
| NEU_AOB_3 | GCA_024281715.1 |
| NEU_AOB_4 | GCA_024281675.1 |
| **JAMMSM_AOB_1** | **GCA_024282015.1** |
| **JAMMSM_AOB_2** | **GCA_024281955.1** |
| **JAMMSM_AOB_3** | **GCA_024281995.1** |
| **JAMMSM_AOB_4** | **GCA_024281935.1** |
| **JAMMSM_AOB_5** | **GCA_024281825.1** |
| **JAMMSM_AOB_6** | **GCA_024281815.1** |
| **JAMMSM_AOB_7** | **GCA_024281795.1** |
| **JAMMSM_AOB_8** | **GCA_024281735.1** |
| **JAMMSM_AOB_9** | **GCA_024281755.1** |
| Pirellula staleyi | GCA_000025185.1 |

| | |
|---|---|
| Candidatus Jettenia caeni | GCA_000296795.1 |
| Candidatus Kuenenia stuttgartiensis | GCA_000315095.1 |
| Candidatus Scalindua brodae isolate RU1 | GCA_000786775.1 |
| Candidatus Brocadia sinica JPN1 | GCA_000949635.1 |
| Candidatus Brocadia fulgida isolate RU1 | GCA_000987375.1 |
| Candidatus Scalindua rubra isolate BSI-1 | GCA_001723765.1 |
| Candidatus Brocadia caroliniensis isolate 26THWARD | GCA_002009475.1 |
| Candidatus Brocadia sp. UTAMX2 | GCA_002050315.1 |
| Candidatus Brocadia sp. UTAMX1 | GCA_002050325.1 |
| Candiadtus Scalindua japonica husup-a2 | GCA_002443295.1 |
| Candidatus Scalindua rubra isolate Ru_enrich_A1 | GCA_002632345.1 |
| Candidatus Kuenenia stuttgartiensis isolate Ru_enrich_A3 | GCA_002632395.1 |
| Candidatus Brocadia sp. R4W10303 | GCA_002848945.1 |
| Candidatus Brocadia sp. isolate AMX2 | GCA_003577195.1 |
| Candidatus Brocadia sp. isolate AMX1 | GCA_003577215.1 |
| Candidatus Brocadia sp. BROELEC01 | GCA_004282735.1 |
| Candidatus Scalindua sp. SCAELEC01 | GCA_004282745.1 |
| Candidatus Scalindua sp. AMX11 | GCA_004351875.1 |
| Candidatus Jettenia ecosi isolate J2 | GCA_005524015.1 |
| Candidatus Brocadia sp. WS118 | GCA_007618135.1 |
| Candidatus Kuenenia stuttgartiensis isolate BL10 | GCA_007618145.1 |
| Candidatus Brocadia sp. BL1 | GCA_007618155.1 |
| Candidatus Brocadia sp. AMX2 | GCA_008363395.1 |
| Candidatus Jettenia sp. AMX1 | GCA_008363445.1 |
| Candidatus Scalindua sp. isolate amx 105 | GCA_008501815.1 |
| Candidatus Scalindua sediminis isolate MohnsGC08_192 | GCA_008636105.1 |
| Candidatus Brocadia sp. isolate SB37 | GCA_008933285.1 |
| Candidatus Scalindua sp. isolate HyVt-343 | GCA_011050115.1 |
| Candidatus Kuenenia stuttgartiensis strain CSTR1 | GCA_011066545.1 |
| Candidatus Kuenenia sp. isolate AMX2 | GCA_011525635.1 |
| Candidatus Brocadia sp. isolate AMX1 | GCA_011525665.1 |
| Candidatus Brocadia sp. isolate AMX3 | GCA_011526495.1 |
| Candidatus Scalindua sp. isolate GLR123 | GCA_013139855.1 |
| Candidatus Brocadia sp. isolate MAG_8 | GCA_013360945.1 |
| Candidatus Brocadia sinica isolate MAG_7 | GCA_013360995.1 |
| Candidatus Brocadia sp. isolate AMX2 | GCA_014338225.1 |
| Candidatus Jettenia sp. isolate AMX1 27 | GCA_014338295.1 |

| | |
|---|---|
| Candidatus Brocadia sp. isolate Ega_18-Q3-R5-49 | GCA_016699285.1 |
| Candidatus Brocadia sinica BroJett002 | GCA_016860665.1 |
| Candidatus Jettenia caeni BroJett041 | GCA_016861385.1 |
| Candidatus Scalindua sediminis isolate Bin_192 | GCA_017368835.1 |
| Candidatus Scalindua arabica isolate SuakinDeep_MAG55_1 | GCA_018263435.1 |
| Candidatus Scalindua sp. isolate SI054_bin45 | GCA_018655105.1 |
| Candidatus Scalindua sp. isolate SI074_bin13 | GCA_018662645.1 |
| Candidatus Scalindua sp. isolate SI075_bin57 | GCA_018669395.1 |
| Candidatus Scalindua sp. isolate SI034_bin70 | GCA_018675875.1 |
| Candidatus Scalindua sp. isolate SI060_bin37 | GCA_018697755.1 |
| Candidatus Scalindua sp. isolate SI072_bin54 | GCA_018700855.1 |
| Candidatus Kuenenia stuttgartiensis isolate BROCD040 | GCA_019454545.1 |
| Candidatus Brocadia sapporoensis isolate BROCD001 | GCA_019455295.1 |
| Candidatus Scalindua rubra isolate MAG_55 | GCA_019911885.1 |
| Candidatus Kuenenia stuttgartiensis isolate MAG_32 | GCA_019912215.1 |
| Candidatus Kuenenia stuttgartiensis isolate FI88_62_9 | GCA_020723505.1 |
| Candidatus Anammoxoglobus sp. isolate DR5_60_7 | GCA_020724075.1 |
| Candidatus Brocadia sp. isolate SJ251 | GCA_020846915.1 |
| Candidatus Brocadia sp. isolate SJ698 | GCA_020851875.1 |
| Candidatus Brocadia sp. isolate SJ692 | GCA_020851925.1 |
| Candidatus Brocadia sp. AMX2 | GCA_021405165.1 |
| Candidatus Jettenia sp. AMX1 | GCA_021405185.1 |
| Candidatus Brocadia sp. isolate DTUAMX1 | GCA_021463075.1 |
| Candidatus Brocadia sp. isolate DTUAMX3 | GCA_021463785.1 |
| Candidatus Brocadia sp. isolate DTUAMX2 | GCA_021463795.1 |
| Candidatus Loosdrechtia aerotolerans isolate YC1 | GCA_021646405.1 |
| Candidatus Brocadia sp. isolate YC2 | GCA_021646415.1 |
| Candidatus Kuenenia stuttgartiensis isolate YC7 | GCA_021646445.1 |
| Candidatus Kuenenia sp. isolate YC6 | GCA_021646465.1 |
| Candidatus Brocadia sp. isolate AM9 | GCA_021650915.1 |
| Candidatus Scalindua sp. isolate OFTM351 | GCA_021776875.1 |
| Candidatus Scalindua sp. isolate OFTM1 | GCA_021776915.1 |
| Candidatus Scalindua sp. isolate OFTM256 | GCA_021776955.1 |
| Candidatus Scalindua sp. isolate OFTM180 | GCA_021777035.1 |
| Candidatus Brocadia sinica HBSIN01 | GCA_022180465.1 |

| | |
|---|---|
| Candidatus Brocadia sapporoensis HBSAPP01 | GCA_022180585.1 |
| Candidatus Jettenia caeni JETCAE04 | GCA_022180965.1 |
| Candidatus Kuenenia stuttgartiensis HKUEN01 | GCA_022180985.1 |
| Candidatus Scalindua sp. SCALA70 | GCA_022181045.1 |
| Candidatus Brocadia sinica isolate MAG.222 | GCA_023150675.1 |
| Candidatus Kuenenia stuttgartiensis isolate AMX1 | GCA_023384235.1 |
| Candidatus Scalindua sp. isolate Bsp_10 | GCA_024223905.1 |
| Candidatus Brocadia sp. isolate AMX2 | GCA_024434125.1 |
| Candidatus Jettenia sp. isolate AMX1 | GCA_024434255.1 |
| Candidatus Kuenenia sp. isolate S40-7 | GCA_024654045.1 |
| Candidatus Scalindua sp. isolate S40-8 | GCA_024654085.1 |
| Candidatus Scalindua sp. isolate P6-7 | GCA_024654465.1 |
| Candidatus Kuenenia stuttgartiensis isolate kuenenia | GCA_900232105.1 |
| Candidatus Kuenenia stuttgartiensis isolate kuenenia | GCA_900232175.1 |
| uncultured Candidatus Kuenenia sp. isolate AMX1 | GCA_900696675.1 |
| Candidatus Loosdrechtia | GCA_21646405.1 |
| Candidatus Brocadia pituitae | GCA_17347445.1 |

**Table S5**: Genome coverage of MAGs in six metagenomes.

| | Genome coverage | | | | | |
|---|---|---|---|---|---|---|
| **Genome** | **Sample 1** | **Sample 2** | **Sample 3** | **Sample 4** | **Sample 5** | **Sample 6** |
| Anammox_bin.281* | 0.9998 | 0.9998 | 0.9998 | 0.9998 | 0.9998 | 0.9998 |
| Anammox_bin.357* | 0.9978 | 0.9978 | 0.9978 | 0.9979 | 0.9978 | 0.9978 |
| JAMMSM_AOB_1 | 0.7252 | 0.7298 | 0.7123 | 0.7354 | 0.7264 | 0.7098 |
| JAMMSM_AOB_2 | 0.6593 | 0.4998 | 0.3596 | 0.4678 | 0.3377 | 0.4389 |
| JAMMSM_AOB_3 | 0.4306 | 0.3308 | 0.3521 | 0.6524 | 0.4410 | 0.5426 |
| JAMMSM_AOB_4 | 0.0000 | 0.0000 | 0.0000 | 0.0000 | 0.0000 | 0.0000 |
| JAMMSM_AOB_5 | 0.0000 | 0.0000 | 0.0000 | 0.0000 | 0.0000 | 0.0000 |
| JAMMSM_AOB_6 | 0.0000 | 0.0000 | 0.0000 | 0.0000 | 0.0000 | 0.0000 |
| JAMMSM_AOB_7 | 0.1161 | 0.0000 | 0.0000 | 0.1327 | 0.0000 | 0.1286 |
| JAMMSM_AOB_8 | 0.0000 | 0.0000 | 0.0000 | 0.0000 | 0.0000 | 0.0000 |
| JAMMSM_AOB_9 | 0.1230 | 0.0000 | 0.0000 | 0.0000 | 0.0000 | 0.0000 |
| JAMMSM_CMX_1 | 0.6658 | 0.7256 | 0.8520 | 0.9112 | 0.7852 | 0.8979 |
| JAMMSM_CMX_2 | 0.3669 | 0.3130 | 0.3586 | 0.4463 | 0.3149 | 0.4111 |
| JAMMSM_NOB_2 | 0.3582 | 0.3523 | 0.3523 | 0.4252 | 0.3325 | 0.3888 |
| JAMMSM_NOB_4 | 0.9994 | 0.9993 | 0.9998 | 0.9999 | 0.9993 | 0.9996 |
| JAMMSM_NOB_5 | 0.3877 | 0.4411 | 0.4367 | 0.5717 | 0.3877 | 0.4747 |
| JAMMSM_NOB_6 | 0.8739 | 0.9049 | 0.9317 | 0.9734 | 0.8899 | 0.9476 |
| JAMMSM_NOB_7 | 0.9271 | 0.9285 | 0.9345 | 0.9664 | 0.9025 | 0.9618 |
| Nitrospira_bin.184* | 0.9993 | 0.9995 | 0.9995 | 0.9996 | 0.9995 | 0.9994 |
| Nitrospira_bin.465* | 0.9976 | 0.9977 | 0.9977 | 0.9978 | 0.9976 | 0.9977 |

* Genomes assembled in this study.

**Table S6:** Relative abundances in in reads per kilobase million (RPKM) of MAGs with average genome coverage higher than 50%.

| | **RPKM** | | | | | |
|---|---|---|---|---|---|---|
| **Genome** | **Sample 1** | **Sample 2** | **Sample 3** | **Sample 4** | **Sample 5** | **Sample 6** |
| Anammox_bin.281 | 15.86 | 14.97 | 16.17 | 14.63 | 13.95 | 15.25 |
| Anammox_bin.357 | 8.77 | 8.29 | 9.04 | 8.23 | 7.83 | 8.43 |
| JAMMSM_AOB_1 | 0.48 | 0.37 | 0.37 | 0.37 | 0.44 | 0.35 |
| JAMMSM_CMX_1 | 0.95 | 0.81 | 0.83 | 1.02 | 0.89 | 0.92 |
| JAMMSM_NOB_4 | 3.87 | 3.97 | 3.94 | 6.30 | 3.66 | 5.44 |
| JAMMSM_NOB_6 | 2.69 | 2.78 | 2.91 | 4.38 | 2.87 | 3.80 |
| JAMMSM_NOB_7 | 2.01 | 1.72 | 1.73 | 2.26 | 1.88 | 2.01 |
| Nitrospira_bin.184 | 3.02 | 3.20 | 3.39 | 5.16 | 3.31 | 4.47 |
| Nitrospira_bin.465 | 3.20 | 3.26 | 3.40 | 5.23 | 3.31 | 4.43 |

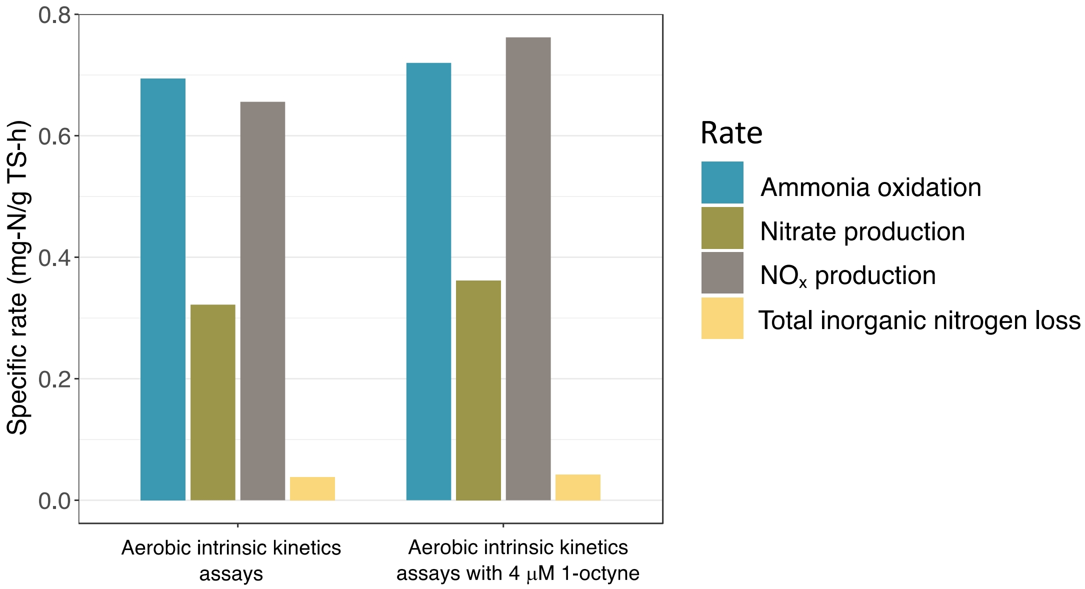


**Figure S1**: Specific rates for ammonia oxidation (blue), NOx production (green), and total inorganic nitrogen loss (yellow) obtained from aerobic ammonia oxidation batch assays for the suspended solids. Bars for each condition are from a single batch assay.

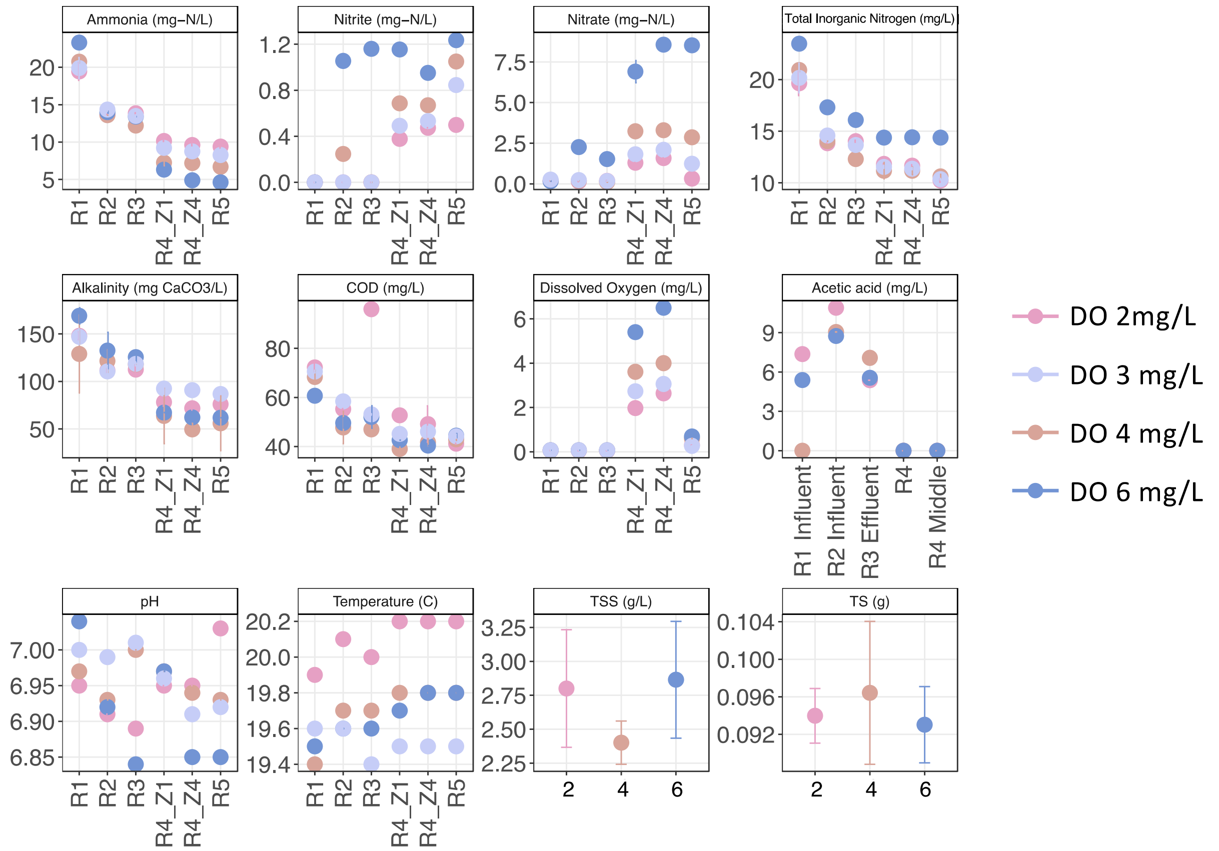


**Figure S2:** Measurements for alkalinity, COD, DO, acetic acid, pH, and temperature sampled along the transect of the IFAS system defined in Section S4 and TSS and TS from the aerobic zone. All samples were taken during full-scale DO set point experiments (2-6 mg/L). Average values were reported for ammonia, nitrite, nitrate, alkalinity, and COD and single values for dissolved oxygen, pH, temperature, and acetic acid. Average values of TSS and TS were reported for the suspended and attached phase. Error bars represent the standard deviations. Influent wastewater contained average ammonia, nitrite and nitrate concentrations of 20.85 (± 1.75), 0 and 0.20 (± 0.04) mg-N/L. Alkalinity, COD, pH, total suspended solids, and total solids measurements were similar in each respective zone for all sampling days. The temperature in the IFAS system was consistent throughout full-scale experiments, ranging from 19.4°C-20.2 °C. Dissolved oxygen concentrations measured in the anoxic zones (R1-R3) were consistently less than 0.07 mg/L, while the desired DO set points in the aerobic zone were achieved for full-scale experiments on their designated days. Slightly higher DO concentrations at the end of the aerobic zone (R4_Z4) were observed compared to the beginning (R4_Z1) but remained within an acceptable range of the DO setpoints. Influent acetic acid concentrations measured on days when the DO set points were 2 and 6 mg/L were 7.37 and 5.39 mg/L, respectively, but were below the LOQ on the day the DO was set to 4 mg/L. Given the detected influent concentrations are close to the LOQ, its possible acetic acid was also present in the influent when the DO was set to 4 mg/L. Across the anoxic zone (R1-R3), acetic acid was produced and then consumed but was not further detected in the aerobic zone.

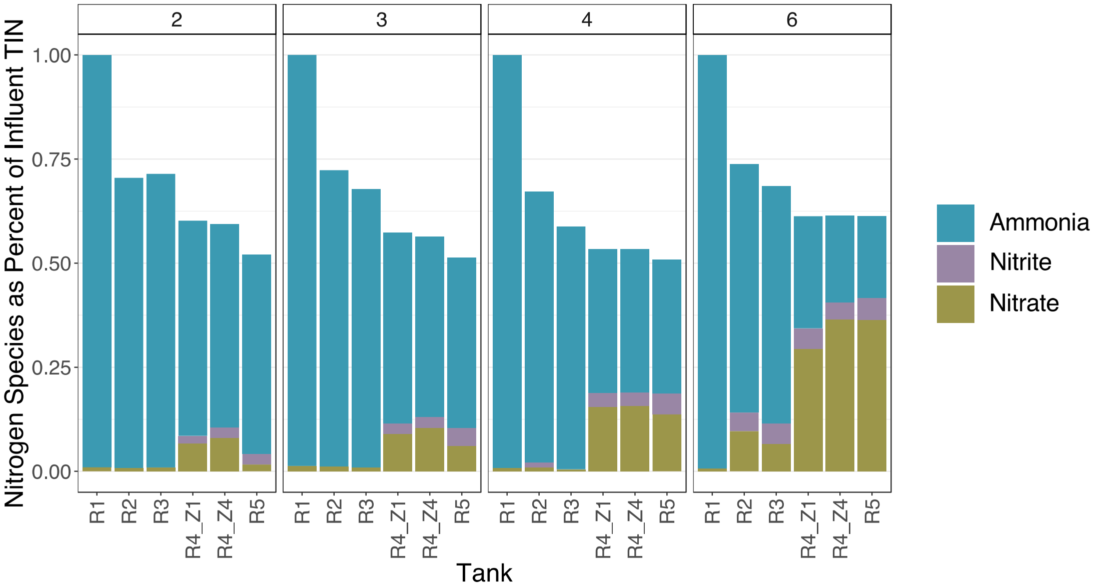


**Figure S3**: Nitrogen species as a percent of influent total inorganic nitrogen (TIN) along the transect of the IFAS system. The facets are labelled based on the DO setpoints. Bars were generated by taking the average concentrations of ammonia (navy), nitrite (orange), and nitrate (green) between sample duplicates and dividing them by the average concentration of influent total inorganic nitrogen. This was done for each sampling location at all DO settings (2-6 mg/L). The average concentration of influent TIN was the summation of ammonia, nitrite, and nitrate concentrations acquired from sample duplicates at R1. Profiles of inorganic nitrogen indicated similar compositions of nitrogen species (mostly as ammonia) entering the aerobic zone when the DO was set to 2, 3, and 4 mg/L, but a higher proportion of nitrite and nitrate entered the zone when the DO was set in 6 mg/L. In this case, nitrate production in the aerobic zone was the highest out of all DO set points, and as a result, more nitrate was returned to the anoxic zone (R2/R3) through the internal recycle line. A dilution effect was observed between R1 and R2 on each sampling day due to the internal recycle coming into the anoxic zone at R2. For example, the average influent ammonia concentration (R1) on one sampling day was 19.45 mg $NH_3$-N/L, and it was diluted to 13.7 mg $NH_3$-N /L in R2.

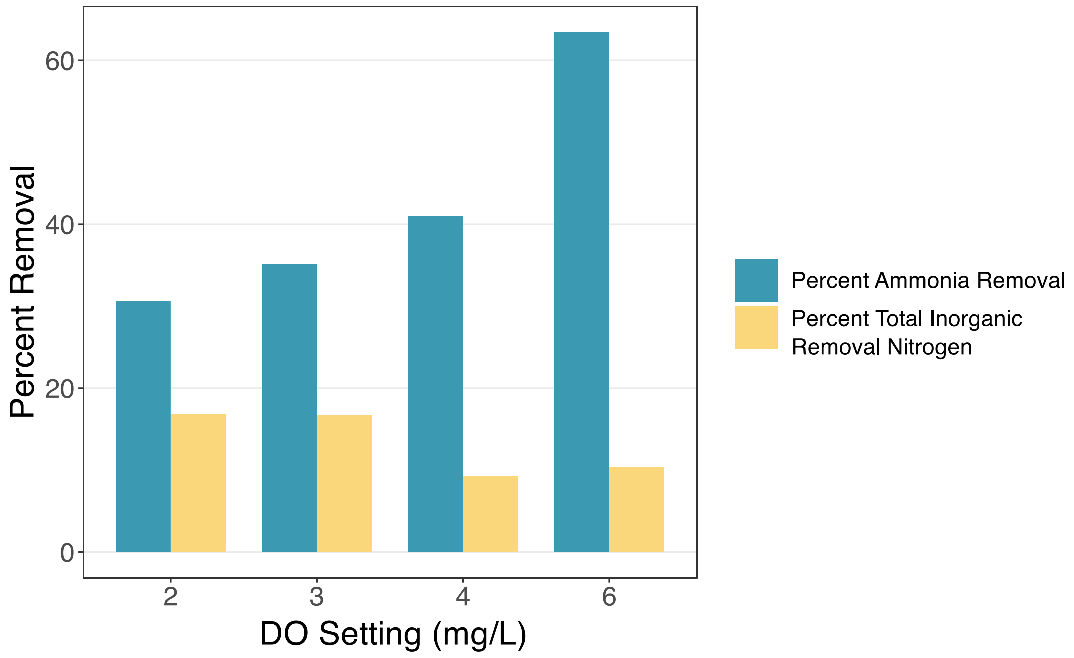


**Figure S4:** Percent of ammonia (navy) and total inorganic nitrogen (yellow) removal in the aerobic zone at each DO set point. Percent removal is calculated from average ammonia or total inorganic nitrogen aerobic zone influent concentration (Figure 1, R3) minus aerobic zone effluent concentration (R4_Z4) divided by the aerobic zone influent concentration.

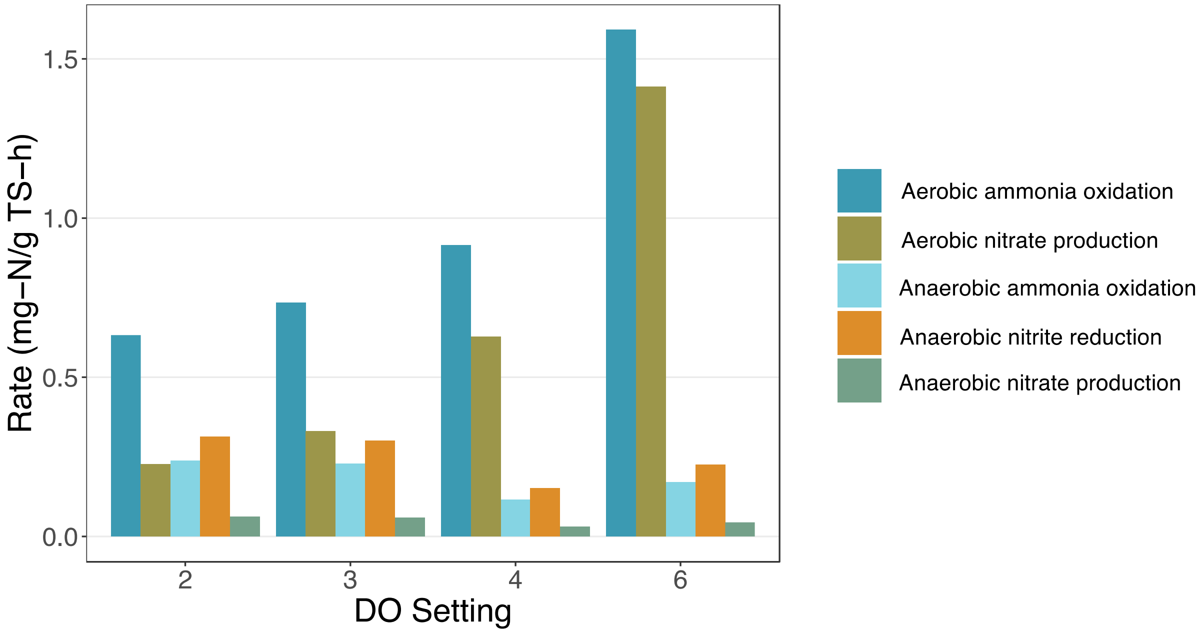


**Figure S5:** Specific rates for aerobic and anaerobic nitrogen biotransformations in the aerobic zone of the full-scale IFAS system during DO set point experiments. Rates were calculated using equations defined in Section S4.
